# Mapping Macaque to Human Cortex with Natural Scene Responses

**DOI:** 10.1101/2025.05.11.653327

**Authors:** Kasper Vinken, Saloni Sharma, Margaret S. Livingstone

## Abstract

Neuroscience has long relied on macaque studies to infer human brain function, yet identifying functionally corresponding brain regions across species and measurement modalities remains a fundamental challenge. This is especially true for higher-order cortex, where functional interpretations are constrained by narrow hypotheses and anatomical landmarks are often non-homologous. We present a data-driven approach for mapping functional correspondence across species using rich, naturalistic stimuli. By directly comparing macaque electrophysiology with human fMRI responses to 700 natural scenes, we identify fine-grained alignment based on response pattern similarity, without relying on predefined tuning concepts or hand-picked stimuli. As a test case, we examine the ventral face patch system, a well-studied but contested domain in cross-species alignment. Our approach resolves a longstanding ambiguity, yielding a correspondence consistent with full-brain anatomical warping but inconsistent with prior studies limited by narrow functional hypotheses. These findings show that natural image-evoked response patterns provide a robust foundation for cross-species functional alignment, supporting scalable comparisons as large-scale primate recordings become more widespread.

## Introduction

Neuroscience depends on cross-species and cross-measurement comparisons to understand the principles of human brain function. Research in humans is mostly based on non-invasive techniques such as functional magnetic resonance imaging (fMRI), which measures broad, indirect signals throughout the entire brain. This technique has been widely used to localize function to cortex, for instance to identify visual regions with response biases for edges, shapes, faces, bodies, or other categories^1–6^. Disentangling the neural computations in these regions requires more focal recording techniques that are largely restricted to animal models. Therefore, much of our understanding of human visual processing has been inferred indirectly from electrophysiological measures in the macaque visual system. A central challenge for systems neuroscience is to connect those findings by aligning responses across measurement modalities and identifying functionally corresponding brain regions across species. Establishing such correspondence is not only critical for interpreting human brain function based on findings from animal models but also provides insights into the evolutionary organization of the cortex^7^.

Existing approaches often rely on narrowly defined functional hypotheses or a small number of diagnostic stimuli to infer correspondence. But such features may fail to capture the full complexity of cortical representations, especially in higher-order areas where conceptual distinctions between regions remain poorly understood^8^. As large-scale, brain-wide recordings in non-human primates become more widely available^9^, and as neuroscience expands into more abstract cortical territories, these limitations become more acute, highlighting the need for scalable, hypothesis-agnostic methods that can leverage such data. Here, we propose one such framework: a flexible, data-driven approach for identifying functionally corresponding brain regions across species using response selectivity to a large number of naturalistic stimuli. By leveraging the rich, graded response patterns evoked by diverse natural scenes, this method identifies functional alignment without relying on predefined tuning axes, stimulus categories, or conceptual heuristics.

To evaluate this approach, we focus on the ventral face patch system, a well-characterized yet still unresolved test case for cross-species alignment. This system comprises a series of patches that respond more to faces than to other objects (“face selectivity”^3,10^). In humans, the ventral face patches form a sequence that extends posteriorly from the occipital face area (OFA) through the fusiform face area’s posterior (FFA-1) and anterior parts (FFA-2) and culminates anteriorly at the anterior temporal lobe face patch (ATL). In the macaque inferotemporal cortex (IT), a comparable sequence unfolds from the posterior lateral face patch (PL) through the middle lateral (ML) and anterior lateral face patches (AL), leading to the anterior medial face patch (AM)^11,12^. These patches contain a high proportion of face-selective neurons called “face cells”, which have been studied extensively in macaque visual cortex^10,13–30^. While both species exhibit a broadly conserved posterior-to-anterior axis, the precise correspondence of specific patches remains unresolved^12^. The macaque central IT face patch ML and human FFA are often considered homologous^11,31–33^. This notion is supported by predictions based on full-brain anatomical warping, which suggests an ML to FFA and an AL to ATL correspondence^33,34^. Nevertheless, evidence from the surrounding cortical topography suggests a possible PL to OFA, ML to FFA-1, and AL to FFA-2 correspondence^35^ or even a PL/ML to OFA and AL to FFA correspondence^36,37^.

To further complicate the picture, anatomical evidence may be insufficient on its own to establish the correspondence of face areas, as regions can reorganize, duplicate, segregate, or enlarge through cortical expansion during evolution^38,39,45^. Functional evidence is therefore essential to determine whether two regions in macaques and humans serve similar roles in visual processing. Traditionally, functional correspondence is determined based on a conceptual interpretation of the role that a region may serve in brain processing. For macaque face regions, such a conceptual distinction is how neurons respond to heads presented at different orientations. Whereas ML neurons show viewpoint-specific responses, more anterior AL neurons show mirror-symmetric viewpoint invariance^16^, a tuning property that can be retrieved from fMRI activation patterns^40^. In humans, one study found evidence for this mirror-symmetric coding in FFA, but not in OFA^41^, suggesting an ML to OFA and AL to FFA functional correspondence. An alternative interpretation is that ML may correspond to the more posterior part FFA-1 and AL to the more anterior part FFA-2^12^. Others have suggested that evidence for mirror-symmetric tuning could merely be explained by low-level confounds^42,43^. The lack of consistency amongst studies calls into question the validity of relying on a single, hand-picked tuning property to arbitrate between scenarios of functional correspondence across species. Indeed, recent findings have shown that, even for face cells, the neural tuning should not be reduced to a few human-interpretable features or concepts^29,30^.

Here, we test whether we can establish the functional correspondence of human and macaque face areas without relying on specific assertions of functional specialization. Rather than focusing on only face images or pre-defined functional properties, we compared neural responses from macaque and human cortex to a shared, diverse set of natural scenes. Specifically, we leveraged an existing large-scale dataset of human fMRI responses to complex natural scenes^44^, and recorded neural responses in macaque central and anterior face regions using the same stimuli. This allowed us to evaluate whether response selectivity across a large number of natural images can resolve competing hypotheses about the alignment of face-selective regions across species. We found that a direct comparison of these responses supported a mapping in which macaque ML corresponds to human FFA and AL to ATL, consistent with predictions from full-brain anatomical warping alone^33^. Thus, our approach resolved prior inconsistencies and demonstrated that broad, natural image-evoked response profiles can establish functional alignment across species, without assuming conceptual distinctions between regions.

## Results

We presented stimuli from the Natural Scenes Dataset^44^ to five macaque monkeys (initials A, OG, P, R, and B1; hereafter referred to as M1-M5), one with an array in primary visual cortex (V1; M1: N=11 reliably responsive multiunit sites, see Methods), two with arrays in CIT at the location of the middle lateral face patch (ML; M2: N=23 units; M3: N=33 units), and two with arrays in AIT targeting the anterior lateral face patch (AL; M4: N=74 units; M5: N=40 units). The stimulus set consisted of a rich set of 700 photographs of a variety of things in a scene context (including animals, humans, sports, food…) captured at varying distances. Most of these stimuli did not prominently feature faces. On average, each of the IT arrays (but not the V1 array) responded most strongly to images with prominent faces (**Fig. 1a**). To evaluate face selectivity of individual units, we computed a face versus no-face d’ metric comparing the response to images with prominent faces to those without faces or animals (see Methods). The units recorded from IT arrays had a high level of face selectivity (mean face versus no-face d’; M2, CIT: 1.46, 95% CI[1.24,1.67]; M3, CIT: 2.12, 95% CI[1.76,2.48]; M4, AIT: 2.30, 95% CI[2.14,2.47]; M5, AIT: 3.71, 95% CI[3.44,3.98]), unlike the V1 array (M1: 0.15, 95% CI[-0.03,0.34]).

**Figure 1.**
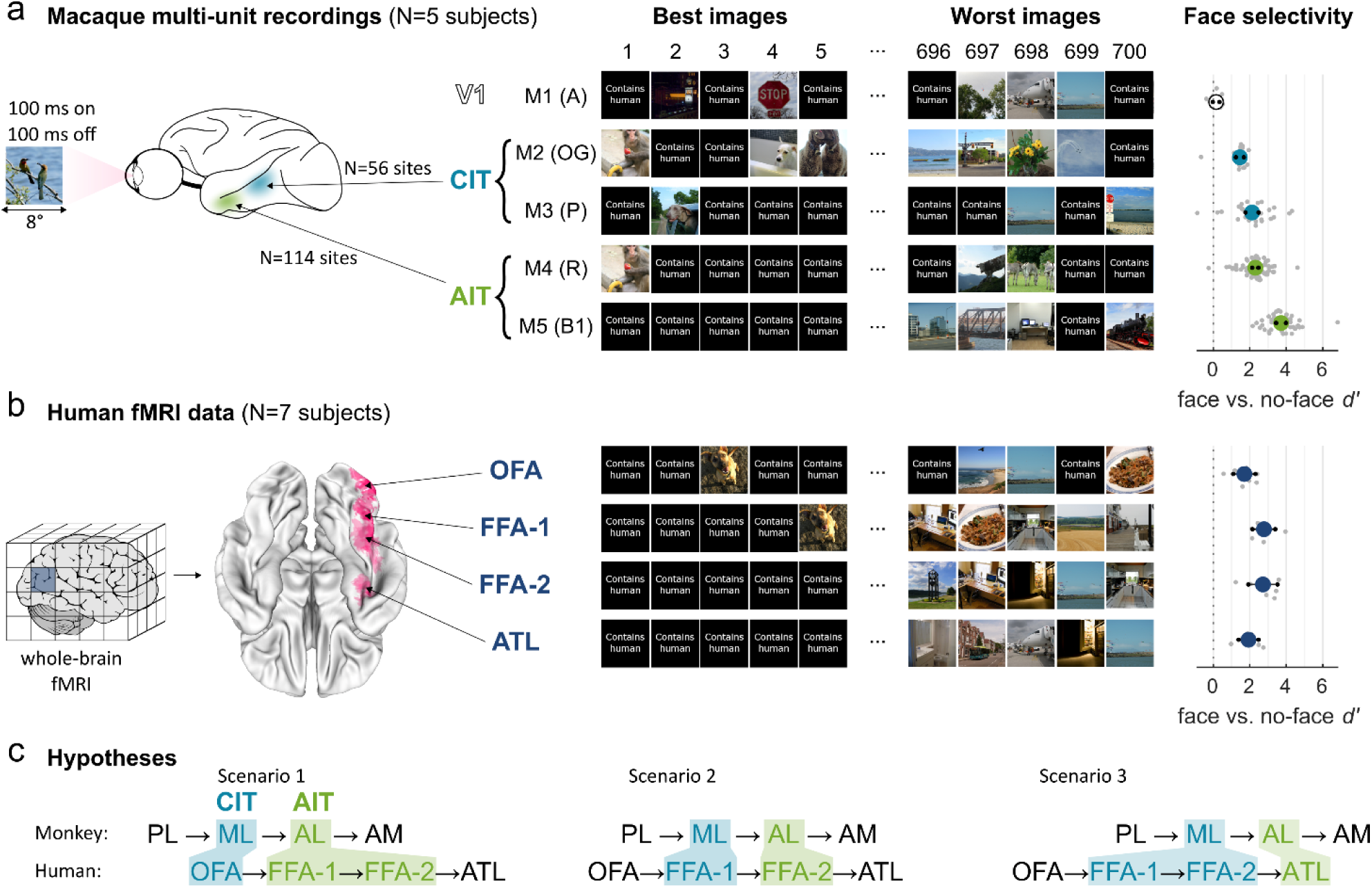
Data and hypotheses. **a.** Central and anterior IT array locations show face-selective responses. Middle: images ranked according to the array response averaged across units. Right: average face selectivity. Grey markers represent individual units. Black markers indicate 95% CI. **b.** Human face-selective ROIs from the functional localizer experiment of ^44^. Middle: images ranked according to the ROI response averaged across voxels and subjects. Right: average ROI face selectivity. Grey markers represent individual subjects. Black markers indicate 95% CI. **c.** Three scenarios of posterior-anterior functional alignment of face-selective cascade in the macaque and human ventral streams.

Similarly, for the human fMRI data, all four localized face area regions of interest (ROIs) responded most strongly to images with prominent faces (**Fig. 1b**). Note that we excluded the mid temporal lobe face ROI, since it was localized in only two subjects. We computed a face versus no-face d’ per ROI of each individual subject, confirming that these regions have strongly face selective responses (mean face versus no-face d’; OFA: 1.70, 95% CI[1.09,2.31]; FFA-1: 2.78, 95% CI[2.15,3.40]; FFA-2: 2.72, 95% CI[1.92,3.52]; ATL: 1.93, 95% CI[1.38,2.47]). Thus, the CIT and AIT arrays were highly face selective, like human face areas, confirming that we successfully targeted the middle and anterior face-selective regions in IT cortex.

To determine whether response patterns across natural scenes can resolve the correspondence between macaque and human face-selective regions, we evaluated three hypothesized alignments (**Fig. 1c**): (1) ML to OFA and AL to FFA; (2) ML to FFA-1 and AL to FFA-2; (3) ML to FFA and AL to ATL. That is, does macaque ML correspond best to human OFA (scenario 1) or to (posterior) human FFA (scenario 2 or 3), and does macaque AL correspond best to human (anterior) FFA (scenario 1 or 2) or to human ATL (scenario 3)? We used array-to-fMRI response pattern similarity to arbitrate between these scenarios.

### Macaque IT arrays correlate with human fMRI face areas and beyond

Having established the face selectivity of the monkey arrays, where in the human brain do we find similar tuning? Rather than characterizing and interpreting the tuning of neural or fMRI responses as a function of specific visual features or categories, we take an agnostic approach where we compare the full selectivity profile across all images. This approach is motivated by recent work from our lab, where we showed that the tuning of face cells is more complex than what can be characterized with faces alone, applying to all kinds of objects in a meaningful way^29,30^. The assumption that we make with this approach is that the stimulus set of 700 images is rich enough to differentiate between distinct complex neural tuning profiles, even if that tuning is too complex to be human interpretable.

We computed the similarity of responses at each specific array location (i.e., V1, CIT, or AIT) to the responses of each vertex of a subject-averaged human brain (**Fig. 2a**; see Methods, Array-to-fMRI similarity). Briefly, for a given vertex in the human brain, we took the trial-then-subject-averaged response vector and calculated the Pearson correlation with each individual unit’s trial-averaged response vector. We also calculated a joint reliability for each unit-to-vertex combination, based on the unit’s trial-wise split-half reliability and the vertex’s subject-wise split half reliability. The macaque array-to-human brain similarity was then computed by averaging for each vertex the Pearson correlations separately across V1, CIT, and AIT units, and normalizing by the noise ceiling given by the unit-averaged joint reliability values.

**Figure 2.**
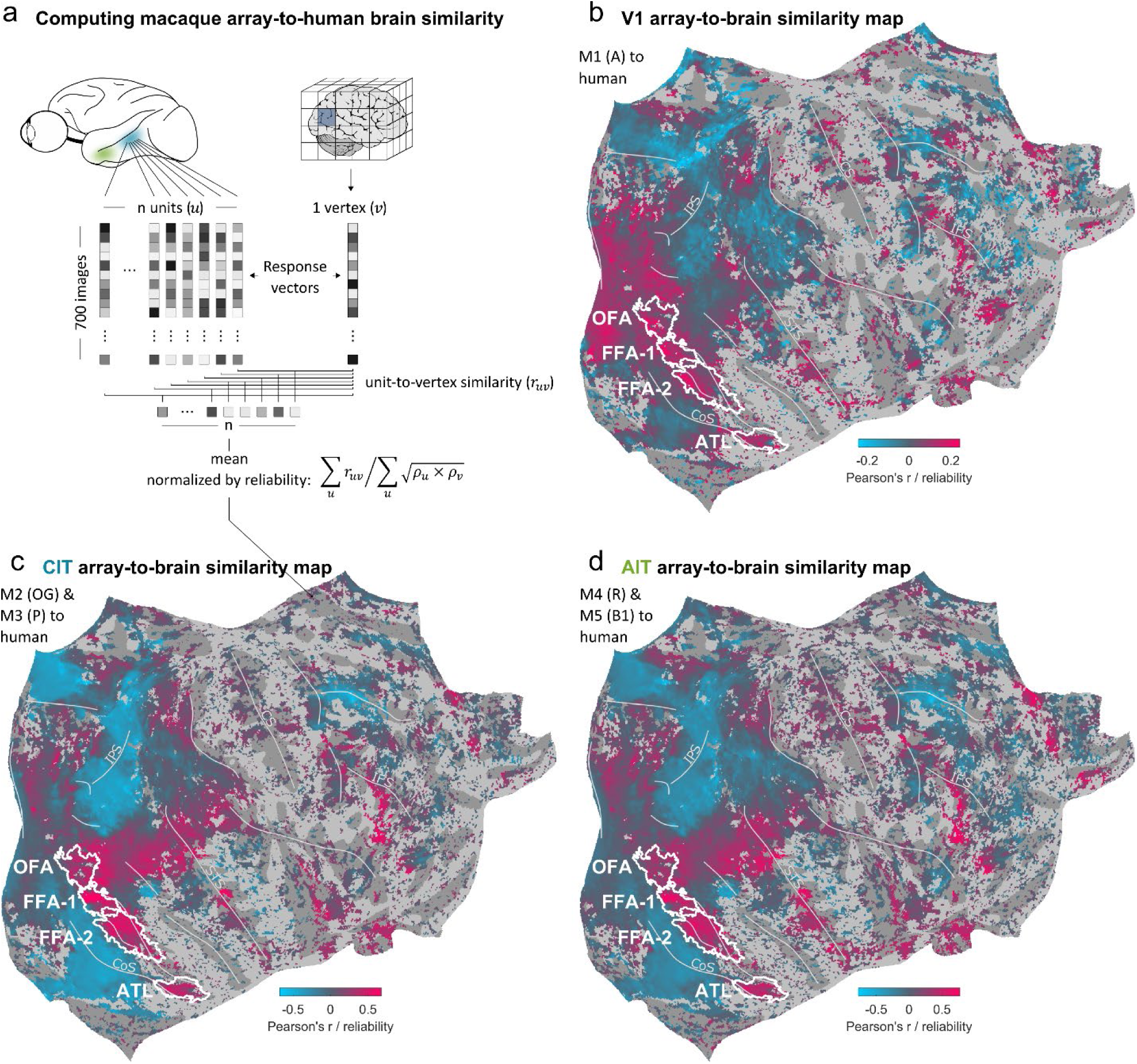
Macaque array-to-human brain similarity maps. **a.** Illustration of how we computed the similarity between array unit responses and human fMRI vertex responses (subject-averaged in fsaverage space). For each vertex *v*, the unit-to-vertex correlations *r_uv_* were averaged across units *u* and then normalized by the Spearman-Brown-corrected joined reliability values 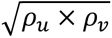, averaged across units *u*. **b-d.** Full right hemisphere human brain maps of similarities with macaque V1 array responses (b), CIT array responses (c), and AIT array responses (d). Vertices for which the noise ceiling (unit-averaged joint unit-vertex reliability) was below 0.2 were masked out. White outlines indicate clusters of vertices for which the fraction of subjects that has a given ROI present exceeds 0.33.

This analysis yielded three macaque array-to-human brain similarity maps, based on the right human hemisphere: one for V1 units (**Fig. 2b**), one for CIT units (**Fig. 2c**), and one for AIT units (**Fig. 2d**). The array-to-brain similarity maps showed a smooth, graded range of negative to positive correlations values, extending well beyond visual areas. This is consistent with a widespread, correlated engagement of the cortex in response to these images. The map based on the macaque V1 array shows similarity biased towards earlier visual areas, although it also shows marked similarity with the higher human face areas. The maps based on macaque CIT and AIT arrays, in contrast, are biased towards higher human visual areas, particularly face regions, as well as regions beyond visual cortex. The perceptual difference between maps based on CIT and AIT was less clear and required a more direct comparison.

To better assess these differences, we Fisher Z-transformed the maps (see Methods, Fisher Z-transformation) and subtracted the map based on CIT from the map based on AIT (**Fig. 3a**). We further masked out vertices that were not correlated or were anticorrelated with both AIT and CIT (Pearson’s *r* / reliability < 0.1). In this AIT minus CIT map, red hues indicate vertices in the human brain that responded more like AIT units than like CIT units. Blue hues indicate vertices that responded more like CIT. This visualization more clearly highlights the fine-grained differences between response similarities with AIT and with CIT. As a sanity check, we subtracted a map based on all IT units from a map based on V1 units (**Fig. 3b**). This V1 minus IT map confirmed that vertices in early visual cortex responded more like V1 units (red hues), whereas vertices in higher visual cortex and beyond responded more like IT units (blue hues).

**Figure 3.**
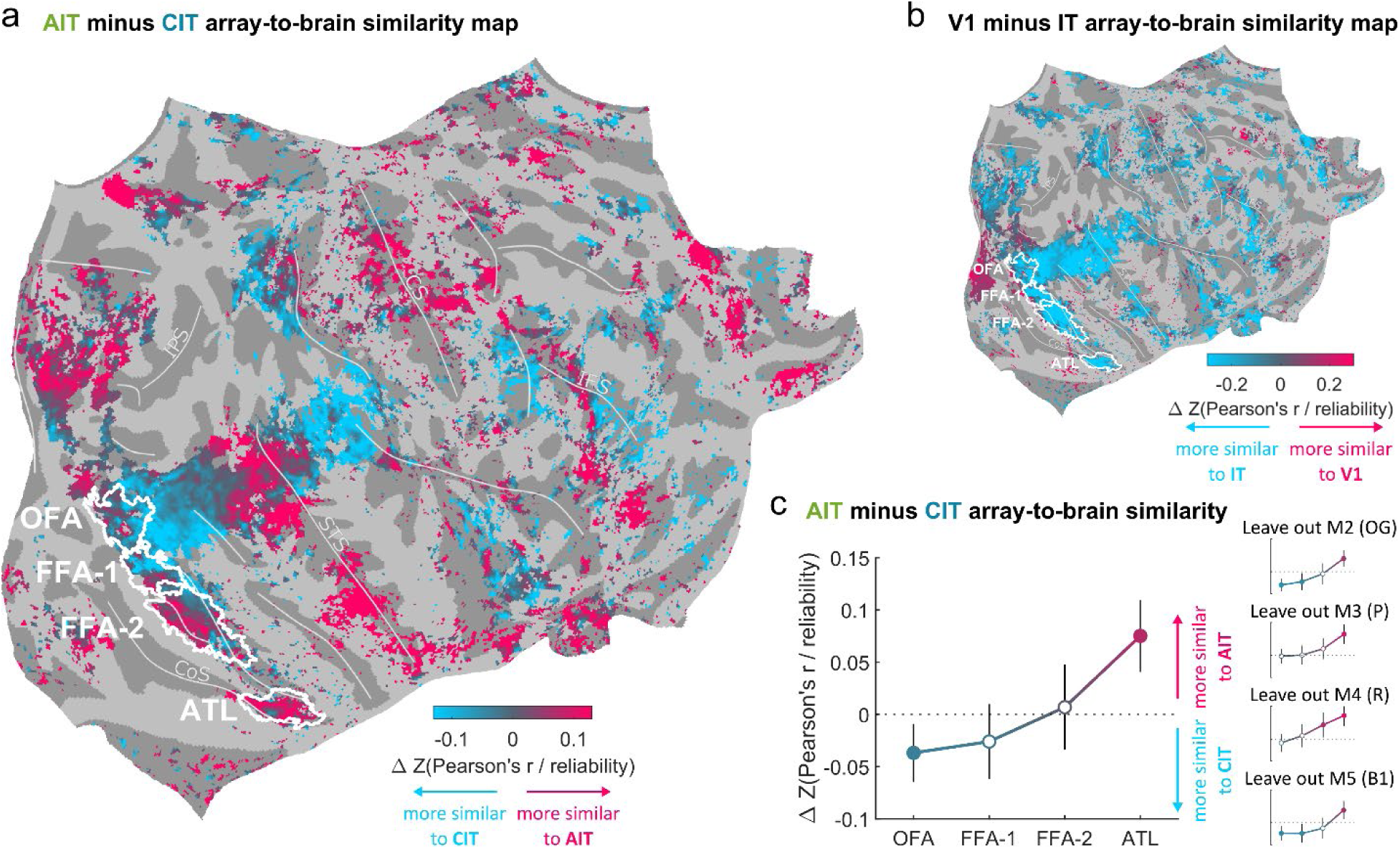
Monkey AIT matched higher human visual (face) areas better than did monkey CIT. **a.** Full right hemisphere human brain map showing the difference in similarity with macaque AIT and CIT array responses (subject-averaged in fsaverage space). Red hues indicate higher similarity to AIT units than to CIT units. Blue hues indicate higher similarity to CIT units than to AIT units. **b.** Human brain map showing the difference in similarity with macaque V1 and IT array responses (red hues: higher similarity to AIT units; blue hues: higher similarity to V1). **c.** AIT minus CIT array-to-brain similarity values averaged for the ROI contours indicated in (a). Error bars indicate 95% CIs based on a two-sample *t*-test comparing AIT units (N=116) to CIT units (N=48). Filled markers denote p<0.05. Insets show the consistency of this trend across monkeys by repeating the analysis while systematically leaving out one monkey at a time (akin to jackknife resampling).

Visual inspection of the response similarities with the target human face regions indicates that human OFA and FFA-1 responses were more like macaque CIT than AIT (predominantly blue hues), whereas human ATL responses were more like macaque AIT than CIT (predominantly red hues). This was less clear for FFA-2: some vertices corresponded better to CIT, others to AIT. For a quantitative analysis, we averaged unit-to-vertex correlations across vertices for which the fraction of subjects that had a given ROI present exceeded 0.33. This analysis confirmed the posterior-to-anterior trend observed through visual inspection, culminating in the largest AIT minus CIT difference in ATL (**Fig. 3c**).

Since FFA vertices did not show a higher similarity to AIT than to CIT units, these results are inconsistent with scenario 1, where the macaque AIT face region AL corresponds to human FFA (and CIT face region ML to human OFA). However, thus far these results remain consistent with both scenario 2 and 3. Next, we used each human subject’s individually localized ROIs in both the left and right hemisphere to determine whether macaque AIT units are better aligned with human FFA-2 (scenario 2) or with ATL (scenario 3).

### Macaque CIT maps best onto human FFA, and AIT onto ATL

To further arbitrate between scenario 2 and 3, and to account for human subject level variability, we computed the similarity of monkey array units to each human subject’s individually defined ROIs. The methods to compute monkey array-to-human brain similarity were analogous to **Fig. 2a** but based on the fMRI voxel responses (trial-averaged beta values) of each human subject’s native space. For each human subject we then averaged array-to-brain similarity values across all voxels within each ROI (see **Fig. 4a**).

**Figure 4.**
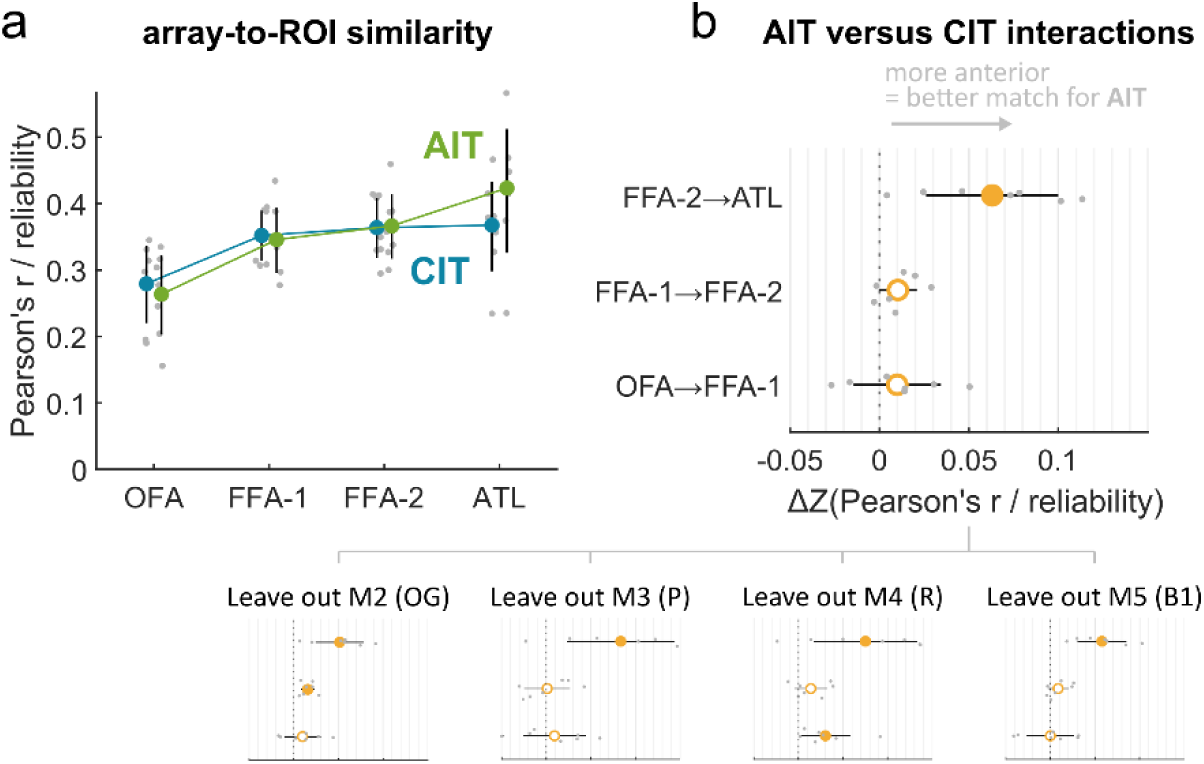
Monkey AIT matched ATL better than it matched FFA. **a.** Average array-to-ROI similarity. Grey markers represent individual human subjects. Error bars indicate 95% CIs based on a one-sample *t*-test (N=7 human subjects). **b.** Regression interaction terms between monkey array (AIT vs. CIT) and a one-step transition in the human ROI cascade (ATL vs. FFA-2, FFA-2 vs. FFA-1, or FFA-1 vs. OFA). For example, for the interaction with ATL vs. FFA-2 (indicated as FFA-2 -> ATL in the figure), the positive value means that the increase in array-to-brain similarity from human FFA-2 to ATL is larger for macaque AIT than for CIT. Error bars indicate 95% CIs based on a paired-sample *t*-test (N=7 human subjects). Filled markers denote p<0.05. Insets show the consistency of this trend across monkeys by repeating the analysis while systematically leaving out one monkey at a time (akin to jackknife resampling).

For both CIT arrays and AIT arrays, there was no significant difference in similarity to human brain responses in FFA-2 versus FFA-1 (CIT: t(6)=0.61, p=0.5639; AIT: t(6)=1.11, p=0.3094). There was also no statistically significant interaction between AIT versus CIT arrays and the difference in similarity to human brain responses in FFA-2 versus FFA-1 (t(6)=2.35, p=0.0571; see **Fig. 4b**). These results are inconsistent with the notion that macaque ML and AL are best distinguished in how well they match human FFA-1 and FFA-2 (scenario 2).

In contrast, there was a significant interaction between AIT versus CIT arrays and the difference in similarity to human brain responses in ATL versus FFA-2 (t(6)=4.17, p=0.0059). Indeed, when pooling similarity to FFA-1 and FFA-2, we found for AIT arrays that similarity to ATL was significantly higher than similarity to FFA (t(6)=2.71, p=0.0353) but not for CIT arrays (t(6)=0.55, p=0.6010). Similarity to OFA was significantly lower than similarity to FFA for both AIT arrays (t(6)=3.81, p=0.0089) and CIT arrays (t(6)=2.66, p=0.0374; confirming the rejection of scenario 1). Thus, overall, these results support the notion that macaque AL is functionally best aligned to human ATL, and macaque ML to human FFA (scenario 3).

## Discussion

We introduce a general framework for identifying fine-grained functional correspondence across species by leveraging population-level neural responses to naturalistic stimuli. Applying this approach to the ventral face patch system, we demonstrate that rich, graded response patterns provide a robust and scalable basis for mapping functional correspondence across species and measurement modalities, without requiring predefined hypotheses or hand-picked stimuli. By directly comparing macaque electrophysiology with human fMRI responses to a shared set of natural scenes, we identified a functional alignment of face-selective areas in which macaque ML corresponds to human FFA and AL to ATL. Despite being derived purely from neural responses, this mapping was consistent with predictions from anatomical topology, suggesting that large-scale cortical layout remains a strong predictor of functional organization across primates. These findings illustrate that neural responses to natural scenes can serve as a reliable basis for cross-species functional inference, offering a flexible, hypothesis-agnostic alternative to traditional methods for comparative neuroscience.

These results help resolve a long-standing ambiguity in the posterior-to-anterior cross-species alignment of face-selective regions. Previous accounts have proposed three competing scenarios: (1) ML aligns with OFA and AL with FFA; (2) ML aligns with FFA-1 and AL with FFA-2; (3) ML aligns with FFA and AL with ATL. Using response patterns across a large set of natural images, we found that recordings from macaque CIT (targeting ML) correlated best with human FFA (both posterior and anterior), with no additional increase for ATL, thereby rejecting scenario 1. In contrast, recordings from macaque AIT (targeting AL) correlated most strongly with human ATL, ruling out scenario 2. Finally, our analyses showed that ML and AL are best distinguished by their correspondence to anterior FFA versus ATL. Together, these findings support scenario 3: a functional alignment of ML with FFA and AL with ATL. This alignment suggests a more anterior human homologue of AL than is often assumed, underscoring the importance of considering broad representational properties in higher-order cortex.

One potential explanation for previous discrepancies based on cortical topography^35–37^ is that evolutionary changes may reshape cortical organization^38,39,45^. Such changes can undermine the reliability of using local topography to infer functional correspondence. On the other hand, our conclusions do converge with a posterior-to-anterior mapping derived from full-brain anatomical warping^33^, suggesting that more holistic anatomical approaches may better capture conserved organizational principles. Previous studies have also compared a range of functional properties, such as mirror-symmetric tuning, the face inversion effect, face familiarity, and face selectivity, often using small sets of diagnostic stimuli^11,12,37,40,46,47^. While these approaches are attractive for their simplicity and interpretability, they are constrained by their reliance on small and highly curated stimulus sets and may miss broader feature tuning. Moreover, recent work has suggested that observations of mirror-symmetric tuning in the human face selective areas may originate from low-level confounds and analysis choices^42,43^. Importantly, these observations highlight a common theme: broader approaches, whether leveraging full-brain anatomical alignment or a comprehensive range of natural stimuli, converge on a consistent cross-species mapping, whereas narrower methods based on local topography or curated stimuli can yield divergent conclusions^48^.

More broadly, our findings illustrate how data-driven approaches using naturalistic stimuli offer a powerful tool for fine-grained functional alignment well beyond the domain of face-selective regions. Traditional methods that rely on small, highly curated stimulus sets risk drawing inaccurate conclusions when the chosen stimuli fail to capture the richness of underlying neural representations. Under stimulus-poor conditions, the inclusion or exclusion of a single image can drastically impact observed response profiles. For example, while face-selective regions in both species responded more strongly to faces on average, the specific images that elicited the strongest responses (Fig. 1, “best” images) differed across species— highlighting nuanced differences in feature tuning that could be either amplified or overlooked with only a few stimuli. These challenges are exacerbated when the stimuli are overly constrained by a conceptual notion such as discrete stimulus categories^49^. Growing evidence suggests that high-level visual cortex, including face-selective cells, is best characterized by an integrated feature space, where category selective responses are carried by domain-general features^29,30,50–53^. By sampling broadly from the natural image space, our approach captures these representational subtleties without imposing categorical assumptions. This strategy enables a more flexible, scalable, and hypothesis-agnostic framework for assessing functional alignment, particularly in higher-order areas where feature tuning is complex or poorly understood.

Establishing cross-species functional correspondence in primates is a long-standing challenge, complicated by issues such as differences in measurement techniques^7^. The advent of fMRI, usable in both monkeys and humans, has enabled an explosion of comparative studies inferring correspondence by comparing fMRI maps of individual functional properties or stimulus contrasts^7,8,11,12,33,35,37–39,46,54–59^. Our approach demonstrates that direct comparisons between electrophysiological and fMRI data can contribute meaningfully to this line of research. By leveraging population-level neural responses and correlating them with human fMRI responses at each vertex, we were able to delineate functional alignment without requiring parallel imaging experiments in both species. Although the physiological differences between spiking and BOLD signals remain, our results show that image-level tuning in multi-unit activity aligns meaningfully with human fMRI responses^60–62^. Indeed, the array-to-brain correlation maps show the highest similarity in regions consistent with each array’s location: human early visual cortex for the macaque V1 array, and ventral temporal cortex (near OFA, FFA and ATL) for the IT arrays. Furthermore, our results converge with previously hypothesized homologies derived from anatomical warping of fMRI-localized macaque face patches onto a human flat map^33,34^. Overall, the fact that our framework captured meaningful alignment across both coarse and fine levels of cortical organization – from early versus high-level visual areas to distinctions among neighboring face regions – suggests that this approach can generalize to systems beyond the ventral stream, wherever shared representations exist across species.

Our study has several limitations. First, we recorded from only one array location per monkey, which means that potential inter-individual differences are confounded with between-area differences. To mitigate this, we repeated the main analyses while systematically leaving out one monkey at a time, confirming that our results are not driven by any single individual. Second, although we focused on two array locations targeting ML and AL, it remains to be seen whether similar cross-species correspondences can be established in other high-level visual areas or cognitive systems. Third, while natural scenes cover a rich and diverse stimulus space, they do not capture the full range of real-world dynamics and behaviors that shape neural activity. Future work could extend this approach by incorporating dynamic stimuli and behavioral context, by sampling simultaneously and more densely from a larger swath of IT or even multiple visual areas^9^, or by applying monkey fMRI to enable full-brain functional warping. Finally, while our approach addresses the question of functional alignment, it does not, by itself, establish phylogenetic homology in the strict evolutionary sense. Such claims require complementary evidence from cytoarchitecture, connectivity, and developmental lineage, which is often inaccessible in humans or for brain regions with ambiguous structural boundaries. In such contexts, our method provides functional evidence to inform hypotheses about homology.

In conclusion, our results point toward a flexible, data-driven framework for establishing functional correspondences across species and modalities. By using naturalistic stimuli to capture rich, high-dimensional and graded response patterns, this approach overcomes the limitations of traditional alignment methods that rely on narrow stimulus sets or predefined functional heuristics. Although we demonstrated this in the ventral visual face patch system, this method is broadly applicable, offering a general framework for establishing cross-species alignment in domains where naturalistic data can be collected. As technological advances continue to enable more large-scale and brain-wide recordings in non-human primates^9^, frameworks that scale with this complexity and support cross-species comparisons of increasingly abstract, hard-to-define functions will be essential. Our findings highlight the value of naturalistic, data-driven alignment as a foundation for such efforts, offering both a principled path toward unified models of brain function and a powerful complement to anatomical and developmental approaches in the study of brain evolution.

## Methods

### NSD data

The human data analyzed in this study were obtained from the Natural Scenes Dataset (NSD), a large-scale fMRI dataset comprising whole-brain, high-resolution measurements collected from eight subjects who viewed natural scenes while engaged in a continuous recognition task. Details regarding data collection and preprocessing can be found in the original publication^44^. Face-selective regions were defined based on the results of the category-selective functional localizer included in the NSD experiments. Labels for these regions were assigned by the NSD authors. For the subject-averaged human brain maps, the provided probabilistic localizer maps were used with a threshold of 0.33 (i.e., only vertices corresponding to a face-selective region in at least one-third of the subjects were included). We additionally defined each subject’s face-selective ROIs individually, based on their native-space face localizer t-maps, using a threshold of t > 3. One subject was excluded because no anterior face region could be defined based on the functional localizer, resulting in a final sample of seven subjects. In addition, a medial temporal lobe face area that could be defined in only two remaining subjects was excluded.

### Animals and arrays

Five adult male macaques were used in this experiment: four rhesus macaques (*Macaca mulatta*; initials A, OG, P, and B1—referred to as M1, M2, M3, and M5, respectively) aged 8-16 years and one pigtailed macaque (*Macaca nemestrina*; initial R, referred to as M4), aged 13 years. All five monkeys were implanted with chronic microelectrode arrays: one in V1 and four in the lower bank of the superior temporal sulcus. Specifically, two monkeys were implanted with 32-channel floating microelectrode arrays (FMA; Microprobes for Life Sciences, Gaithersburg, MD): V1 of M1 and CIT of M3. Three monkeys were implanted with 64-channel NiCr microwire bundle arrays (Microprobes for Life Sciences, Gaithersburg, MD)^63^: CIT of M2 and AIT of M4 and M5. The target location for the face patch arrays was identified using fMRI face localizers (M3, M4, M5) or anatomical landmarks (M2; STS ‘bumps’: Arcaro et al^64^). All procedures were approved by the Harvard Medical School Institutional Animal Care and Use Committee and conformed to National Institutes of Health guidelines provided in the Guide for the Care and Use of Laboratory Animals.

## Experiments

During recordings, the monkeys were performing a fixation task in which they were rewarded with drops of juice to fix their gaze on a spot in the middle of a 53-cm LCD monitor. The gaze position was monitored using an ISCAN system (ISCAN, Woburn, MA). The experiments were controlled with MonkeyLogic (https://monkeylogic.nimh.nih.gov/). During fixation, images were presented at a size of 8 visual degrees and a rate of 100 ms on and 100 ms off. The images were presented at the center of the mapped receptive field, with a jitter of ±1 to ±2 visual degrees for IT arrays. An average of 32 trials were presented per stimulus.

### Stimuli

The stimulus set consisted of a subset of 1000 images that were shared across human subjects in the NSD dataset^44^. The NSD image set contains a rich variety of photographs taken from Microsoft’s Common Objects in Context (MS COCO^65^) image database. For this paper, we focus on 700 images for which there were at least two presentations for each human subject, to be able to compute a split-half reliability using the exact same images for each subject.

### fMRI-guided array targeting

The target location for the array placement for 3 out of 4 monkeys (M3, M4, M5) was identified using an fMRI face localizer. The details of the fMRI experiments are described in Arcaro and Livingstone^66^ and will only be summarized here briefly. The monkeys were scanned using custom-made four-channel surface coils (by A. Maryam at the Martinos Imaging Center), in a 3T TIM Trio scanner with an AC88 gradient insert. Functional images were acquired using a repetition time = 2s, echo time = 13ms, flip angle = 72°, iPAT = 2, matrix size = 96 x 96 mm, resolution = 1-mm isotropic, and 67 contiguous sagittal slices. To enhance signal-to-noise ratio and increase contrast ^67^, the monkeys were injected with 12 mg/kg of monocrystalline iron oxide nanoparticles (Feraheme, AMAG Pharmaceuticals, Cambridge, MA, USA) before each scanning session. The face localizer consisted of randomly shuffled 20s blocks of face or nonface objects, interleaved with 20s of a neutral gray screen.

### Data analysis

#### Firing rates

To compute average firing rates, we counted the number of spikes in a 200 ms window following stimulus presentation onset. The latency of the response was selected based on a visual assessment of the global average (across channels and stimuli) peristimulus time histogram, ranging from 65 ms to 125 ms.

Following previous procedures^29,30^, we used an a-priori response reliability criterion of >0.4 to include only visually driven, selective neural units for further analysis. This yielded 11 multiunit sites from V1 recordings, 48 multiunit sites from CIT recordings and 116 multiunit sites from AIT recordings.

#### Response reliability

We determined a trial-wise split-half reliability per neural unit and per voxel in each NSD subject brain, and we determined a subject-wise split-half reliability per vertex in the NSD ‘fsaverage’ space. To obtain trial-wise split-half reliability, we first computed the correlation 𝑟𝑟 between the average response vector based on odd trials and the one based on even trials. We then applied the Spearman-Brown correction to obtain a reliability 𝜌 = 2𝑟/(1 + 𝑟).

To obtain subject-wise split-half reliability, 𝑟𝑟 was computed as the correlation obtained from the trial-and-subject averaged response vectors for every possible way to split the NSD subjects in two, averaged across splits.

#### Face selectivity

We assessed face selectivity by computing the face versus no-face dʹ sensitivity index comparing trial-averaged responses to a subset of images with prominent primate faces (N=26 out of all 700 images) versus all images without faces or animals (N=255 out of all 700 images):

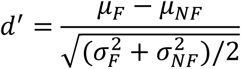

where 𝜇_𝐹_ and 𝜇_𝑁𝐹_ are the across-stimulus means for faces and non-faces, and 𝜎_𝐹_ and 𝜎_𝑁𝐹_ are the across-stimulus SDs.

#### Array-to-fMRI similarity

We quantified the similarity between microelectrode array responses and fMRI vertex/voxel responses as follows. For each fMRI vertex/voxel *v* and array multiunit 𝑢, we computed the Pearson correlation 𝑟*_uv_* between their respective response vectors across images. For the flat-maps, we then averaged the unit-to-vertex correlation values across all array units 𝑢𝑢 to obtain a single correlation value for each vertex *v*. For the ROIs, we averaged across all units and voxels to obtain a single correlation value for each ROI.

To estimate a noise ceiling, we computed joint reliability values as 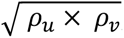 , where 𝜌_𝑢_ and 𝜌_v_ represent the response reliability of each unit and each vertex/voxel, respectively (see Methods, Response reliability). Similar to the correlation values, we averaged these joint reliability values across units (and voxels in the case of ROIs) to obtain a single noise ceiling value for each vertex *v* or each ROI.

Finally, we normalized each array-to-vertex/ROI correlation by its corresponding noise ceiling to obtain a single array-to-vertex/ROI similarity value.

### Fisher Z-transformation

The correlation-based array-to-fMRI similarity values are inherently bounded between -1 and 1 and exhibit a skewed distribution, especially near the extremes. Therefore, for statistical comparison between similarity values associated with a given vertex/ROI, we applied the Fisher Z-transformation, 𝑍(𝑟) = atanh(𝑟), to normalize the distribution and stabilize variance across the range of similarity values^68^.

## Funding

This work was supported by the Alice and Joseph Brooks Fund Postdoctoral Fellow (K.V.), Gordon Fellowship (to S.S.) and NIH grant R01 EY025670 (to M.S.L.)

## Author contributions

Conceptualization: K.V. and M.S.L. Data curation: K.V. Formal analysis: K.V. Methodology: K.V. Investigation: S.S. and M.S.L. Visualization: K.V. Supervision: M.S.L. Writing—original draft: K.V. and S.S. Writing—review and editing: K.V., S.S. and M.S.L.

## Competing interests

The authors declare no competing interests.

## Notes

### Competing Interest Statement

The authors have declared no competing interest.

